# Two decades of suspect evidence for adaptive DNA-sequence evolution – Less negative selection misconstrued as positive selection

**DOI:** 10.1101/2020.04.21.049973

**Authors:** Qipian Chen, Ziwen He, Xiao Feng, Hao Yang, Suhua Shi, Chung-I Wu

**Affiliations:** State Key Laboratory of Biocontrol, Guangdong Key Lab of Plant Resources, Key Laboratory of Biodiversity Dynamics and Conservation of Guangdong Higher Education Institutes, School of Life Sciences, Sun Yat-Sen University, Guangdong, China; CAS Key Laboratory of Genome Sciences and Information, Beijing Institute of Genomics, Chinese Academy of Sciences, Beijing, China; Department of Ecology and Evolution, University of Chicago, Chicago, Illinois, USA

**Author notes:** These authors contributed equally to this work (Qipian Chen, Ziwen He). Correspondence should be addressed to C.-I.W. or S.S.

## Abstract

Evidence for biological adaptation is often obtained by studying DNA sequence evolution. Since the analyses are affected by both positive and negative selection, studies usually assume constant negative selection in the time span of interest. For this reason, hundreds of studies that conclude adaptive evolution might have reported false signals caused by relaxed negative selection. We test this suspicion two ways. First, we analyze the fluctuation in population size, N, during evolution. For example, the evolutionary rate in the primate phylogeny could vary by as much as 2000 fold due to the variation in N alone. Second, we measure the variation in negative selection directly by analyzing the polymorphism data from four taxa (*Drosophila, Arabidopsis*, primates, and birds, with 64 species in total). The strength of negative selection, as measured by the ratio of nonsynonymous/synonymous polymorphisms, fluctuates strongly and at multiple time scales. The two approaches suggest that the variation in the strength of negative selection may be responsible for the bulk of the reported adaptive genome evolution in the last two decades. This study corroborates the recent report^1^ on the inconsistent patterns of adaptive genome evolution. Finally, we discuss the path forward in detecting adaptive sequence evolution.

## Introduction

The rate of molecular evolution is accelerated by positive selection, and retarded by negative selection, relative to the baseline neutral mutation rate. A central task in molecular evolutionary studies is to estimate the amount of positive selection from the evolutionary rate of DNA sequences. Hence, the task would entail sorting out the two forces.

The rate, usually expressed as the ratio between the nonsynonymous (Ka or dN) and synonymous (Ks or dS) changes, is a relative measure independent of time. It can be expressed, in a most reduced form, as

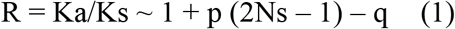

where p and q are the proportion of advantageous and deleterious nonsynonymous mutations, respectively^1–6^, N is the effective population size of a haploid species and s is the selective advantage^7,8^. The fixation probability of all deleterious mutations is assumed to be ~0. Slightly deleterious mutations that have a non-trivial probability of being fixed will require an extension of Eq. (1).

A survey of published sequences found that genome average values of R range between 0.1 and 0.3 across a wide array of taxa including vertebrates, invertebrates and plants^5^. Using Eq. (1), we can see that q is 0.7 at the minimum, thus indicating massive eliminations by negative selection. Obviously, positive and negative selection cannot be simultaneously inferred from R. For example, R = 0.2 may mean p = 0 and q = 0.8 but it can also mean p = 0.01, 2Ns = 11 and q = 0.9. Since p = 0.01 indicates extensive adaptive DNA evolution, the difference between the two cases is biologically highly significant. In this study, we separately analyze the effects of positive and negative selection by comparing DNA sequences both within and between species. The analyses raise serious questions about the adaptive DNA sequence evolution reported in the literature of the last two decades^9–16^.

## PART I Negative selection in relation to population size

### Theory

Various assumptions have been made in the literature to sort out factors affecting positive and negative selection in sequence evolution^17–21^. Among these factors, the most obvious one would be the variation in population size, N. Given that these assumptions are often difficult to validate, we may, instead, ask whether methods based on different assumptions would give the same results on positive selection. The most commonly used tests are the MK test (the McDonald and Kreitman test^18,20^) and the PAML test (Phylogenetic Analysis by Maximum Likelihood^22,23^). In a companion paper^1^, the two commonly used methods are found to yield nearly non-overlapping results, thus raising questions about the validity of some, or even all, of these assumptions.

In the test of positive selection, the null model assumes no positive selection (p = 0) and Eq. (1) is reduced to

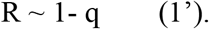

The null model further assumes constant negative selection, i.e. constant q, on the phylogeny of interest. By Eq. (1), if R deviates from constancy, positive selection would be implicated. While a constant q seems reasonable, R would remain constant only when all deleterious mutations are lost. Since weakly deleterious mutations, which account for the bulk of mutations^21,24,25^, can be fixed with a non-trivial probability^5,26^, a generalized form of Eq. (1) is shown by Eq. (2)

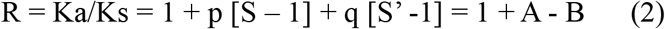

where
and f(N, s) = (1-e^−2s^) / (1-e^−2Ns^) is the fixation probability of advantageous mutation with s being the selective advantage of the mutation^3,7,8^.

In testing DNA sequence evolution, we set p = 0 in the null model and the equation of interest is reduced to

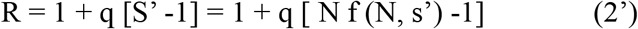

where f(N, s’) = (1-e^2s^’) / (1-e^2Ns^’) is the fixation probability of deleterious mutations with s’ being the selective strength^4,8,27^. Eq. (2’) is reduced to Eq. (1’) if f (N, s’) = 0, when deleterious mutations are never fixed. Condition for f (N, s’) > 0 are given in Fig. 3 of Chen *et al*. (2019)^5^.

By Eq. (2’), the assumption of constant negative selection is in effect a declaration that the effective population size on the phylogeny of interest stays constant. Such a declaration is difficult to justify and, indeed, genomic studies of various taxa have documented substantial changes in N^28–33^.

### Hypothesis testing based on empirical data

Eqs. (2) and (2’) lead to two postulates on selection efficacy and population size.

1. The efficacy of selection, in particular negative selection, may not stay constant due to the changes in N across lineages and time intervals.
2. The efficacies of positive and negative selection are likely positively correlated if the variation in N plays a large role in Eq. (2’).

To test the first postulate, we estimate the historical effective population sizes of three primates using the PSMC model (Pairwise Sequentially Markovian Coalescent model)^28^. It can be seen in Fig. 1a-c that N fluctuates substantially in the last 5 Myrs (million years). During this time span, N fluctuates by 15, 7 and 12-fold in humans, chimpanzees and macaques, respectively. To see how this fluctuation may affect the Ka/Ks ratio, we plot the fate of deleterious mutations that have a fitness effect of s = −1/(2N*) where N* is the lowest N in the last 5 Myrs in macaques (see Methods for details). In this setting, the physiological effects of mutations (in terms of q and s) are assumed the same among primates.

**Figure 1.**
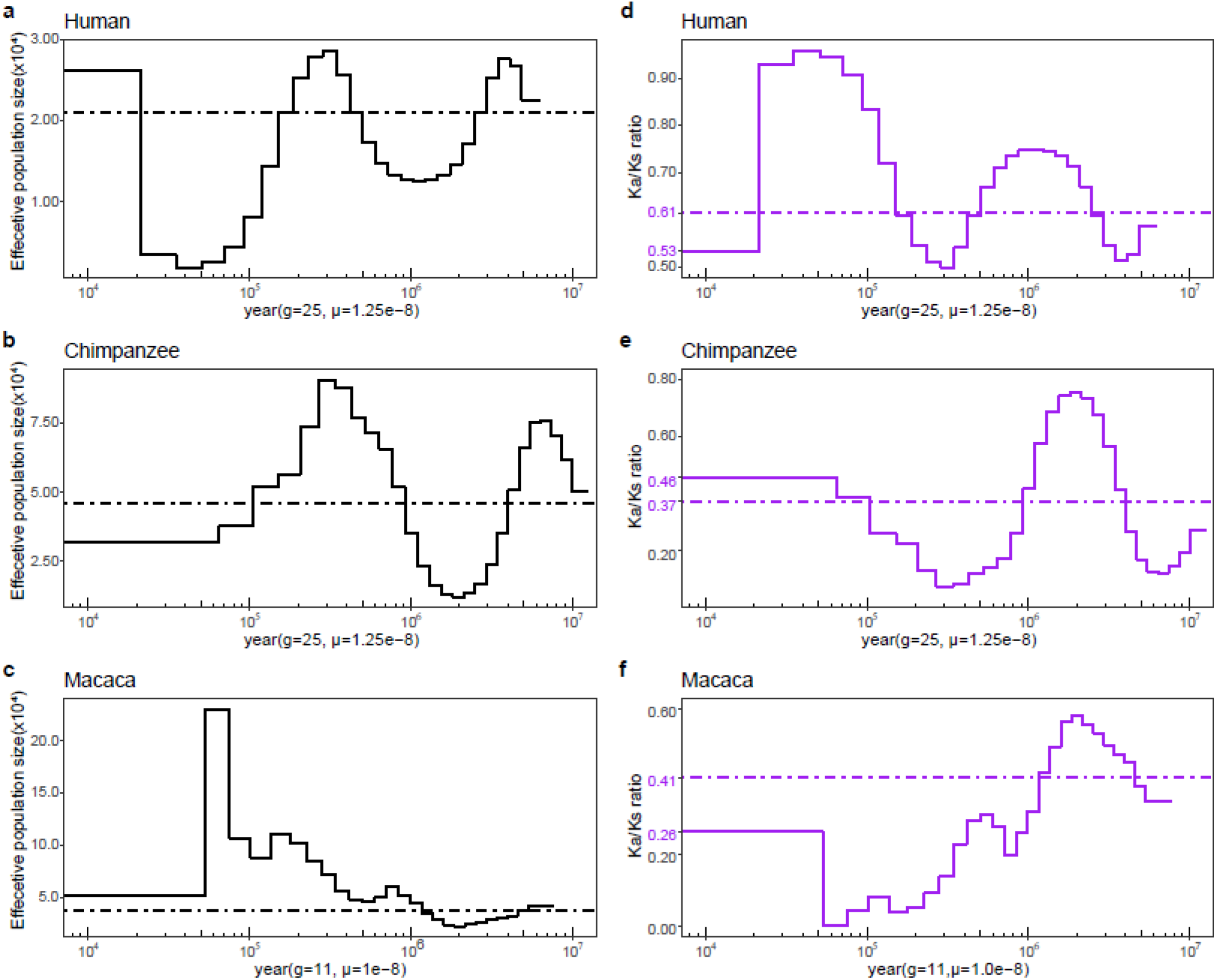
Temporal effective population size and Ka/Ks ratio. **a-c**, inferred historical population sizes from pairwise sequential Markovian coalescent analyses in humans, chimpanzees, and macaques, respectively. **d-f**, changes to Ka/Ks ratios over time according to population size, assuming all nonsynonymous mutations are deleterious (s=-2.18×10^−5^, see Methods). Dash lines represent average values over the past 5 million years.

Fig. 1d-f show the variations in Ka/Ks around the broken line that depicts the average in this time span, at 0.61, 0.37 and 0.41 for humans, chimpanzees and macaques, respectively. For these weakly deleterious mutations, the Ka/Ks ratio fluctuates wildly in the last 5 Myrs around a different mean for each species. Even between humans and chimpanzees, the strength of negative selection has been quite divergent. Across the 3 lineages, the lowest Ka/Ks ratio is 0.00042 in the macaque lineage and the highest one is 0.95 in humans. This > 2000 fold difference may have happened between 30 and 60 Kyrs before present (see Fig. 1d and f).

Another noteworthy value is the Ka/Ks ratio in the most recent past, which has been the proxy of the evolutionary average in the MK test^34^. We note that the extant ratio is larger than the historical value by 50% in chimpanzees (0.46 vs. 0.37) whereas the trend is opposite in macaques (0.26 vs. 0.41). Overall, the changes in N in the last 5 Myrs appear to contradict the common assumption of constant negative selection over this time span.

## PART II Direct measurement of negative selection from the polymorphism data

### Theory

While the fluctuation in N can be a major factor governing negative selection, there is a threshold for Ns’ above which the fluctuation may not matter. For example, whether |Ns’| = 100 or 1000, S’ is essentially 0. The value of |Ns’| matters the most when it is close to, or smaller than, 1^4,5,35^. Since we know little about the distribution of Ns’, it would be necessary to estimate the strength of negative selection directly from the polymorphism data of each species. We designate Pa and Ps for the polymorphism measure of nonsynonymous and synonymous changes, analogous to the Ka and Ks distance between species. Pa/Ps = 1 is the neutral ratio and Pa/Ps < 1 is indicative of negative selection.

A complication is that Pa/Ps depends on the frequency of the variants (x) in the population. Fig. 2a shows the frequency spectrum for variants under different intensities of selection, ranging from Ns’ = −10 to Ns = 50. Fig. 2b shows the spectrum under negative selection relative to the neutral ones. These spectra can be equated with those for the Pa/Ps ratio, if one assumes synonymous variants are neutral (but see Lu and Wu, 2005^36^). Pa/Ps decreases as the frequency increases since slightly deleterious mutations can often reach a detectable frequency before being eliminated. Nevertheless, the rate of decrease is strongly dependent on the Ns’ value (Fig. 2b). Fig. 2c shows the effect of positive selection, resulting in an increase in the high frequency portion of the spectrum (x > 0.75). The trends in Fig. 2b and Fig. 2c are opposite, as expected.

**Figure 2.**
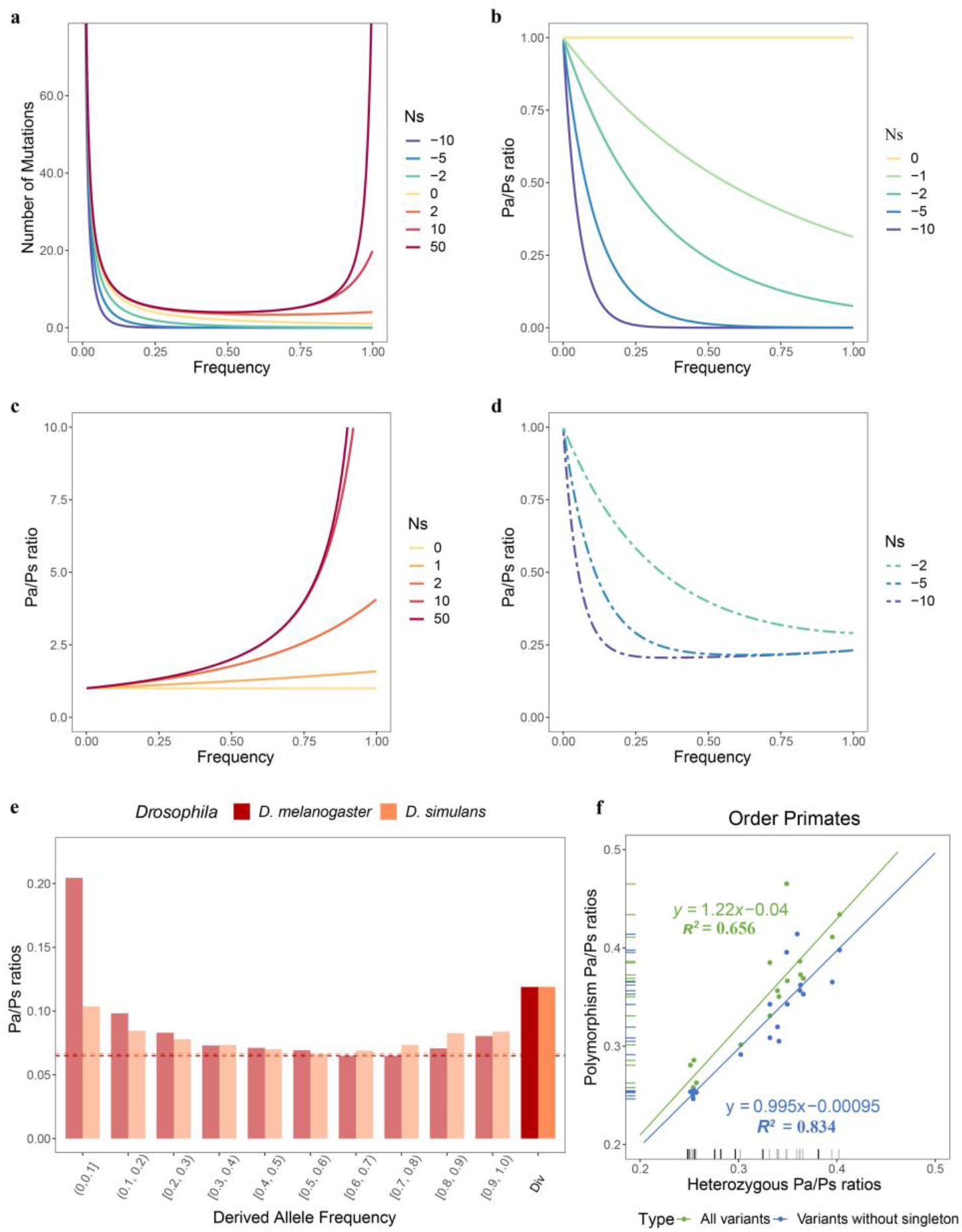
Frequency spectrum of nonsynonymous mutations in populations. **a**, Frequency spectra of mutations under different selection strength. **b-d**, the Pa/Ps ratios as a function of frequency, where all synonymous mutations are assumed to be neutral. Nonsynonymous mutations in each curve in b-c is either under negative (a), positive(c), or no selection (b and b). Dashed curves in (d) depict nonsynonymous mutations are under both negative (q=0.8) and positive selection (p=0.01, Ns=2). **e**, Pa/Ps ratios in *D. melanogaster* and *D. simulans*, binned by frequency of derived variants. The opaque bars depict the lineage specific fixed Pa/Ps ratios, which are equivalent to the Ka/Ks ratio. **f**, Two levels pf Pa/Ps ratios of non-CpG sites in Primates. The rugs at the X axis represent heterozygous Pa/Ps ratios of primate species with only one individual. The grey lines represent the species which have many individuals.

In reality, the spectrum would be a mixture of mutations under positive and negative selection. In Fig. 2d, the composite spectra with p = 0.01 and q = 0.8 are shown for various strengths of negative selection (Ns = −2, −5 and −10). The effect of negative selection is reflected in the portion of the spectrum that is “flat” (i.e., most deleterious mutations having been eliminated). It is also prudent to avoid variants of the very high frequency, which could be beneficial mutations that have not been fixed. If Ns < −5, the variants suitable for estimating negative selection should be those with x between 0.25 and 0.9 (Fig. 2d). If Ns is weaker, e.g. Ns = −2, the estimation of negative selection would be much more difficult.

Fig. 2e shows the Pa/Ps spectrum in *Drosophila* which yields an estimate of Pa/Ps ~ 0.07 after deleterious mutations are filtered out (x > 0.2). As the bulk of mutations are in the range of x < 0.2 (see Fig. 1a), the sample size has to be quite large to yield the “flat” portion of the spectrum. For this reason, it is not possible to accurately estimate the strength of negative selection in taxa where extensive polymorphism data are not available. In this study, we remove only the singleton class, i.e., variants that appear once in the sample. Although this practice may not remove all deleterious mutations, it is informative about the relative strength of negative selection among species.

### Additional source of empirical data – Genomes from one individual

In many species, there is no data on polymorphism. Nevertheless, the complete sequences of a diploid individual consist of two haploid genomes, which could provide information on polymorphism^30,37–41^. If genomes of single individuals can be used in lieu of a polymorphism survey, then the database would expand greatly. We now use the polymorphism surveys from primates to gauge the feasibility of using one single individual to estimate the Pa/Ps ratio.

In primates, 17 species/subspecies have been subjected to population genomic sequencing. For these species, the Pa/Ps ratio can be computed at either the population or the individual level. The computation of the population level Pa/Ps is described above for the 5420 orthologs available across species. At the individual level, we compute the ratio of nonsynonymous/synonymous heterozygosity (per site) for each individual and take the average across individuals. In this treatment, the correlation between the individual-level and population-level Pa/Ps ratio is high with R^2^=0.83 (see Fig. 2f and Table 1). In short, heterozygous SNPs of each individual are informative and many more species with genomic data from one single individual can be used for investigating the variation in negative selection.

**Table 1.** Summary of the polymorphism patterns in primates.

| Species (subspecies or population) | N<sup>a</sup> | $\theta\pi$ (per kb) | Pa/Ps ratio | | | Ka/Ks<sup>c</sup> |
| --- | --- | --- | --- | --- | --- | --- |
|  |  |  | Average heterozygosity<sup>b</sup> | Polymorphism (all sites) | Polymorphism (no singletons) |  |
| *Homo* |  |  |  |  |  |  |
| *H. sapiens* (Han Chinese) | 106 | 0.278 | 0.390±0.022 | 0.520 | 0.400 |  |
| *H. sapiens* (Yoruba population) | 107 | 0.397 | 0.378±0.015 | 0.449 | 0.382 | 0.329±0.013 |
| *Pan* |  |  |  |  |  |  |
| *P. paniscus* | 13 | 0.440 | 0.437±0.024 | 0.419 | 0.384 | 0.321±0.013 |
| *P. troglodytes verus* | 4 | 0.430 | 0.429±0.017 | 0.397 | 0.353 |  |
| *P. troglodytes ellioti* | 10 | 0.786 | 0.393±0.014 | 0.373 | 0.344 |  |
| *P. troglodytes schweinfurthii* | 6 | 0.874 | 0.368±0.016 | 0.344 | 0.309 | 0.327±0.013 |
| *P. troglodytes troglodytes* | 4 | 0.964 | 0.370±0.011 | 0.338 | 0.295 |  |
| *Gorilla* |  |  |  |  |  |  |
| *G. beringei graueri* | 3 | 0.400 | 0.397±0.022 | 0.356 | 0.341 | - |
| *G. gorilla gorilla* | 23 | 0.964 | 0.360±0.011 | 0.372 | 0.331 | 0.325±0.013 |
| *Pongo* |  |  |  |  |  |  |
| *P. pygmaeus* | 5 | 0.902 | 0.360±0.009 | 0.320 | 0.298 | - |
| *P. abellii* | 5 | 1.246 | 0.328±0.010 | 0.291 | 0.282 | 0.302±0.013 |
| *Rhinopithecus* |  |  |  |  |  |  |
| *R. roxellana* | 18 | 0.412 | 0.380±0.028 | 0.355 | 0.332 | 0.301±0.017 |
| *R. bieti* | 20 | 0.253 | 0.393±0.009 | 0.359 | 0.349 | 0.338±0.016 |
| *Macaca* |  |  |  |  |  |  |
| *M. mulatta lasiotus* | 32 | 1.347 | 0.276±0.009 | 0.275 | 0.245 |  |
| *M. mulatta littoralis* | 29 | 1.305 | 0.272±0.007 | 0.270 | 0.244 |  |
| *M. mulatta tcheliensis* | 5 | 1.106 | 0.275±0.011 | 0.244 | 0.240 | 0.310±0.018 |
| *M. mulatta breviceaudatus* | 5 | 1.121 | 0.275±0.014 | 0.248 | 0.237 |  |
| *M. mulatta mulatta* | 10 | 1.297 | 0.278±0.005 | 0.253 | 0.243 |  |

<sup>a</sup> N, sample size of each species (subspecies or population).
<sup>b</sup> Average heterozygosity Pa/Ps ratios with standard deviation in N individuals of each species or subspecies.
<sup>c</sup> Average Ka/Ks ratios with standard deviation compared to each species in other families. (Between Hominoids and old-world monkeys).

We should note that the calibration of Fig. 2f is applicable only to primates. Each taxon, say, *Drosophila*, will need its own calibration. Nevertheless, if the focus is only on the variation in the strength of negative selection, then the genomic data from one individual per species are still informative. Naturally, one would only know whether the strength varies among species without knowing the precise strength.

### Hypothesis testing based on empirical data

We now use the polymorphism data (including one pair of genomes from a single individual) to investigate the variation in the strength of negative selection among extant species. The taxa are *Drosophila* (4 species), *Arabidopsis* (4 species or sub-species), primates (17 species) and birds (38 species). These data cover plants, invertebrates and vertebrates.

#### 1) Drosophila

The four *Drosophila* species shown in Fig. 3a are taxa commonly used for probing adaptive molecular evolution^11,14,15,20^. Clearly, the selective constraint fluctuates wildly even in this small group. Most notable is *D. sechellia*, which has a much higher Pa/Ps value than others. Among the rest, Pa/Ps at low frequency (<0.2) is higher in *D. melanogaster* than in *D. simulans* and *D. yakuba*, but above 0.2, the Pa/Ps values are similar (0.062-0.065, see Fig. 3a and Table S1). To see the trend, we plot the Pa/Ps ratio of each species in Fig. 3b against its level of polymorphism, the latter reflecting the effective population size^8^. There is a clear negative correlation suggesting that the larger the effective population size, the stronger the selective constraint. This negative correlation is consistent across taxa as the rest of this study shows.

**Figure 3.**
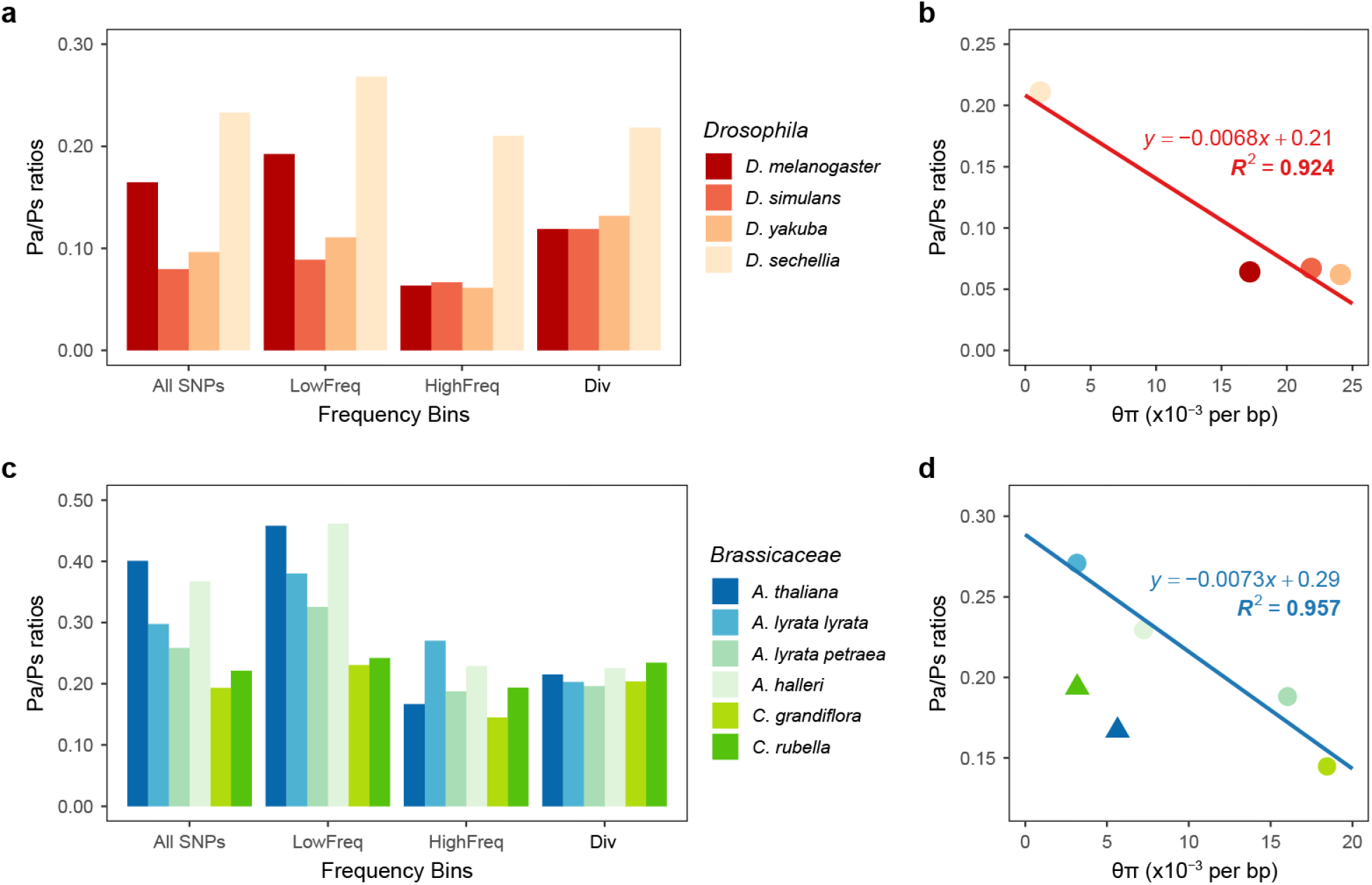
Non-synonymous to synonymous ratio between and within species. **a** and **c**, the Pa/Ps ratios grouped by frequency and Ka/Ks ratios in *Drosophila* and *Brassicaceae*. Variants with frequency below 0.20 are grouped into low-Freq bins and other polymorphisms are grouped into high-Freq bins. The bars at Fixed bins represent lineage-specific Ka/Ks ratios, labeled on the branches of the corresponding color on the phylogenetic trees in Fig. S1 and Fig. S2. **b** and **d**, Neutral genetic diversity (θπ of synonymous polymorphism) and Pa/Ps ratios in *Drosophila* and *Brassicaceae*. Triangles represent self-compatible species.

The patterns raise some interesting issues. If one wishes to use the MK test to detect positive selection among the four species, one would compare the Pa/Ps ratio(s) with the Ka/Ks value between species. Fig. 3a shows that the lineage-specific Ka/Ks ratios are comparable, ranging between 0.120-0.200. Hence, the comparison between the interspecific Ka/Ks and the polymorphic Pa/Ps of *D. sechellia* would show no evidence of adaptive evolution. Among the remaining three species, the results are more nuanced. If all variants are used (as is commonly done), the conclusion for adaptive evolution would depend on the source of the polymorphism data – negative when using the data from *D. melanogaster* but positive when using the data of from the other two species. This is why Fay *et al*. (2002) proposed to impose a cutoff of gene frequency^42^. Many other procedures have since been introduced^11,43,44^. When one uses polymorphisms with the mutant frequency at > 0.2 (see Fig. 3a), all the remaining species yield signals of positive selection.

Obviously, the main difficulty in detecting positive selection is that negative selection is not constant. The non-constancy of negative selection also violates the assumption of PAML. However, the robustness of PAML in detecting positive selection may require more extensive analyses of past changes in Ne (such as those done in Fig. 2). In *Drosophila*, the MK and PAML tests show very marginal overlaps in the adaptive signals detected^1^. Because of the differences in estimating negative selection, they may have both interpreted the relaxation of negative selection as positive selection, albeit in different manners.

#### 2) Arabidopsis

*Arabidopsis thaliana* and its relatives are the main model organisms among plants, with high-quality reference genomes and polymorphism data^45,46^. Here, we investigate their selective constraints. The divergence of *A. thaliana* and *A. lyrata* is 11%, similar to the divergence between *D. melanogaster* and *D. simulans* (10.5%). However, negative selection in *Arabidopsis* fluctuates more wildly than in *Drosophila*. With the low frequency SNPs (|0.2) removed, Pa/Ps is substantially higher in *A. lyrata* (*subsp. lyrata*) and *A. halleri* than in *A. thaliana* and *A. lyrata* (*subsp petraea*) (see Fig. 3c), whereas their lineage-specific Ka/Ks ratios are similar, ranging from 0.198 to 0.225.

Following the common pattern, each species’ Pa/Ps ratio is a function of the effective population size, which is reflected in the diversity measure (θ; see Fig. 3d)^8^. In this comparison, the correlation is given for the outcrossing species only. The self-compatible species of *A. thaliana* and *Capsella rubella* (triangles in Fig. 3d) have much lower Pa/Ps ratios than other species with a comparable diversity (θ). This trend is unsurprising as self-compatible species generate homozygotes at a high rate even for deleterious mutations that are rare. The results raise the question on the MK test again. When using the two taxa with the lower Pa/Ps (*A. thaliana* and *A. petraea*), one would conclude positive selection. But, using the ratios from the other two species (*A. lyrata* and *A. hallerĩ*), one would reach the opposite conclusion.

Compounding the issue, the polymorphism patterns are rather different among these species and the cut-off to filter out the low-frequency variants should be different for each species (Fig. 3c and Table S2). *Arabidopsis* is therefore a typical example that the variation in the strength of negative selection, as manifested in Pa/Ps, is so large that the detection of positive selection based solely on DNA sequence data would be unreliable.

#### 3) Primates

For primates, we compile the data from 17 species (plus one subspecies) belonging in 6 genera. Since hominoids and old-world monkeys (OWMs) have diverged by < 6% in their DNA sequences, these two families are very close in molecular terms. This level of divergence would be suitable for detecting positive selection by both the MK and PAML test. (For a comparison, the two morphologically identical sibling species of *D. melanogaster* and *D. simulans* have diverged by more than 10%.)

The polymorphism and divergence data are presented in Table 1 and Fig. 4. The polymorphism Pa/Ps ratio again varies considerably among species. The trend appears to be an increasing Pa/Ps ratio toward human. With singleton and CpG sites removed, the polymorphism Pa/Ps ratio in primates decreases in the following order: human and bonobo (0.400-0.384), chimpanzee and gorilla (0.353-0.295), orangutan (0.298-0.282) and macaque monkeys (0.245-0.237) (see Table 1 and Table S3). The snub-nosed monkey (0.349-0.332), an old-world monkey, is the only group that deviates from the general trend.

**Figure 4.**
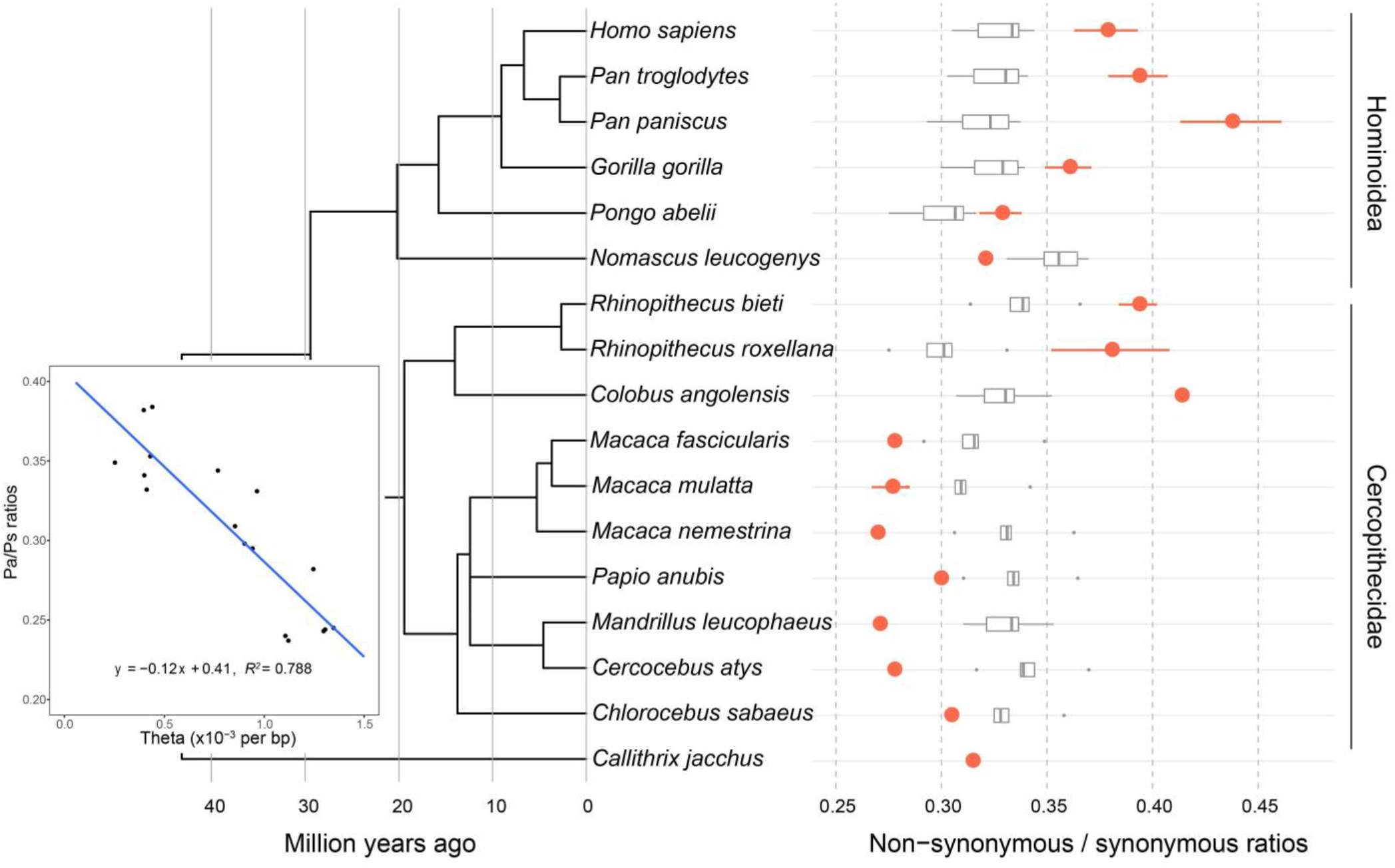
Non-synonymous to synonymous ratios in primates. The grey boxplots at right show the Ka/Ks ratios of species across families. The red points and attached error bars show the mean and standard deviations of heterozygous Pa/Ps ratios. The scatterplot shows the correlation of neutral genetic diversity (θπ estimated from synonymous polymorphism) with Pa/Ps ratios.

As shown in the inset of Fig. 4, the Pa/Ps ratio is again negatively correlated with the diversity estimate of θπ, which is a function of the effective population size. The R^2^ value is 0.79, corroborating the predictions of Eqs. (1) and (2). In short, a gradual reduction in the population size from old world monkeys to apes and humans may have played a large role in the step-wise relaxation of negative selection in this direction.

The trend of a larger Pa/Ps ratio toward apes and human casts doubts on the inferences of adaptive evolution in the genomic sequences of primates. Note that Ka/Ks > Pa/Ps is often assumed to be the hallmark of positive selection. The Ka/Ks ratio for each lineage, say human (any ape species), is the comparison with every OWM species, thus yielding a distribution as shown. Likewise, each OWM species is compared with every ape. If we compare human and any OWM species, the Ka/Ks value is 0.31 – 0.34. Since the Pa/Ps is human is about 0.38, there is no evidence of adaptive evolution between human and OWMs. However, since Pa/Ps ranges between 0.26 and 0.30 among OWMs, one would conclude positive selection between the same two taxa.

The contradiction just stated is very general when one compares any ape species with any OWM species. In Fig. 4, the top half (including all apes, *Rhinopithecus* and *Colobus*) shows that the Pa/Ps ratios (red dots) generally fall in the higher range of 0.32 – 0.44 while the 7 OWM species in 5 genera of the lower half have Pa/Ps between 0.25 and 0.32. Importantly, the Ka/Ks ratio generally falls between the two sets of Pa/Ps values. In short, the assumption of constant negative selection is violated between apes and OWMs, thus precluding the inference of adaptive evolution in genomic sequences between them.

#### 4) Birds

For birds, we analyze the genomic data of 38 species from 30 orders. The general trend is the same as those of the three taxa shown in Figs. 3 and 4. In Fig. 5, the Pa/Ps ratios are scattered between 0.131 and 0.266 among all bird species with a mean of 0.179 and standard deviation of 0.035 (see Table S4). The Pa/Ps ratio is again negatively correlated with diversity (measured by the average heterozygosity; see the inset of Fig. 5) with R^2^ = 0.593. In contrast, the Ka/Ks ratio falls in a relatively small range of 0.152 −0.235. Again, the variation in the strength of negative selection in birds, as seen in the Pa/Ps value relative to the interspecific divergence in Ka/Ks, is far too large to permit the analysis of positive selection based solely on the genomic sequences.

**Figure 5.**
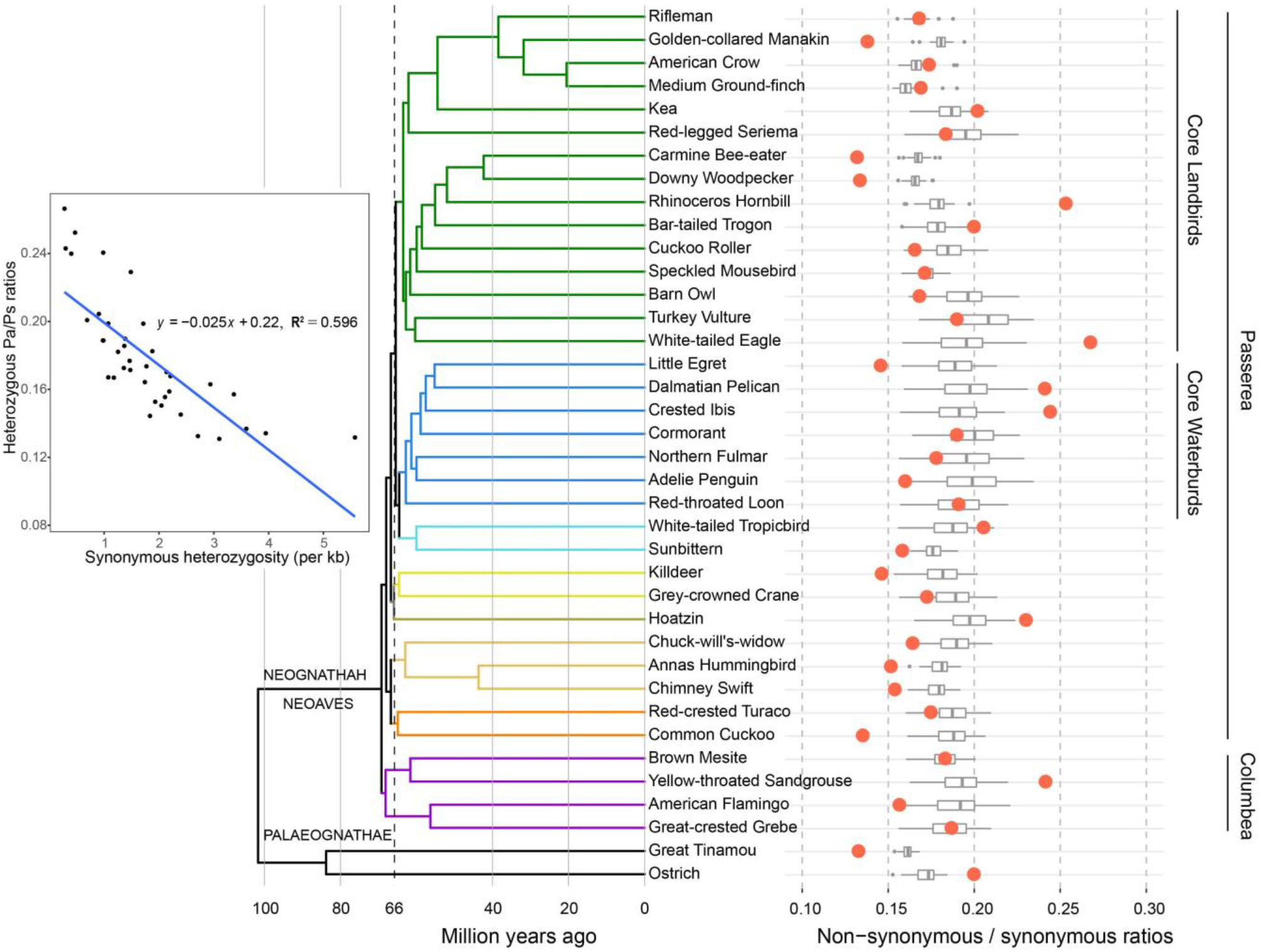
Non-synonymous to synonymous ratios in birds. The tree at left was modified from the highly resolved total evidence nucleotide tree inferred with ExaML by Jarvis *et al*. (2015)^39^. The colors of branches denote well-supported clades. The red points in the right panel represent heterozygous Pa/Ps ratio of each species and the grey boxplots represent distributions of Ka/Ks ratios of divergence from other species on the tree. The scatter plot shows the negative correlation of neutral heterozygosity (synonymous heterozygosity) with Pa/Ps ratios.

## Discussion

For two decades, the search for signals of adaptive sequence evolution has been done at the expense of understanding negative selection. An expedient assumption that negative selection is constant in the time frame of interest is often made. With this assumption, any deviation from the expected rate of DNA evolution must be due to positive selection. Because different tests (mainly MK and PAML) would interpret different aspects of negative selection as adaptive signals, they may reach divergent conclusions, both likely false, on adaptive evolution. Indeed, He *et al*. (2018) have shown that the two tests often yield non-overlapping results^1^.

In our analyses, negative selection is found to deviate strongly from the assumed constancy in all taxa tested. Hence, much of the reported adaptive evolution in DNA sequences in the last two decades is likely the consequence of variation in negative selection, rather than the operation of positive selection. A central claim of the Neutral Theory of molecular evolution that DNA sequence evolution is shaped mainly by genetic drift and negative selection still stands, 50 years after its proposal^47^.

The variation in negative selection has been obvious to evolutionists. First, the population size changes in multiple time scales and across lineages; hence, the fixation probability of deleterious mutations would fluctuate across branches in any phylogeny. Second, if negative selection is constant, then based on Eq. (1)

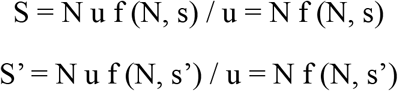

In other words, the observations of Ka/Ks < Pa/Ps, as seen frequently in Figs. 4 and 5, contradict the constancy assumption. Given p ≥ 0, q on the two sides of the ≥ sign cannot possibly be the same one. A more accurate version should be Eq. (2) but the argument is the same.

Whether the genomic sequences carry information on past adaptation remains a central issue in molecular evolutionary studies. It is clear that sufficient efforts in quantifying negative selection must accompany the detection of positive selection. Incorporating population-size fluctuation, as done in Fig. 1 using the PSMC method, could be a first-step. Nevertheless, a method that can directly estimate the changes in the strength of negative selection, rather than indirectly estimate the effective population size, may be more desirable in the long run.

Finally, the strengths of positive selection and negative selection are strongly correlated because, as shown in Eq. (2), both are function of N. The correlation may in fact be more general than Eq. (2) shows. For example, large-step mutations, when measured by the physico-chemical distances between amino acids, are both more deleterious and more beneficial than small-step mutations^26,48^. Therefore, the working of negative selection would be informative about the operation of positive selection as well. In conclusion, to understand adaptive evolution at the DNA level, we must analyze positive and negative selection concurrently and in the same context.

## Methods

### Divergence Data

#### 1) Genus Drosophila

Coding sequences (CDS and their translated proteins) and *D. melanogaster* based orthologs were downloaded from FlyBase (http://www.flybase.org)^49,50^. We used the FB2019_03 gene models. Protein sequences were aligned using default parameters in MUSCLE v3.8.31^51^. We then aligned CDS of orthologous genes with the help of aligned protein sequences using PAL2NAL v14^52^. We obtained 9679 alignments of the *melanogaster* subgroup species (*D. melanogaster, D. simulans, D. sechellia, D. yakuba*, and *D. erecta*) spanning 11 million years of evolution (see Fig. S1)^14,53,54^. Genes with high divergence rates, apparently caused by misalignment, were discarded. Only genes with more than 100 codons were used. The final dataset contains 7042 orthologous autosomal gene sets (2L, 2R, 3L, and 3R).

#### 2) Family Brassicaceae

We obtained the longest transcripts sequences of protein-coding genes from *Arabidopsis thaliana, Arabidopsis lyrata, Arabidopsis halleri, Capsella grandiflora*, *Capsella rubella*, and *Boechera stricta* from the PhytozomeV12 database^55–60^. Pairwise orthologs for the six species were inferred using the InParanoid algorithm^61^. Only genes with a single 1:1 ortholog for each species were used. We obtained protein alignments using MUSCLE v3.8.31^51^. And we obtained CDS alignments using aligned protein sequences and PAL2NAL v14^52^. We began with 13737 alignments of the six species and filtered the data as for *Drosophila*. The final dataset consists of 11735 orthologous genes.

#### 3) Order Primates

In the order Primates, 16 Catarrhini species have genome assemblies in NCBI (The National Center for Biotechnology Information)^62–74^. We used the common marmoset genome data as the outgroup. A summary of assemblies and annotations of these 17 species is found in Table S5. The phylogenetic tree was downloaded from the Time Tree website (http://www.timetree.org/). Using the software OrthoFinder v2.2.7^75^, optimized with Diamond^76^, we identified 7538 autosomal *Homo sapiens*-centric orthologous gene sets of 17 primate species. CDS and protein alignments were obtained using MUSCLE v3.8.31 and PAL2NAL v14^51,52^. Because mammals exhibit hyper neutral CpG mutation rate^77,78^, we masked codons with CpGs to TpG/CpAs changes.

#### 4) Class Aves

The CDS and translated sequences of protein coding genes of 38 birds were obtained from the Avian Phylogenomic Project website^38,39,79^. We used OrthoFinder v2.27 with the aid of Diamond to find 8295 orthologs^75,76^. The MUSCLE-PANLNAL pipeline described above was used to align protein coding sequences. Raw reads for genome assembly were downloaded from NCBI to obtain polymorphisms from single individuals. A summary of genome assemblies and raw reads is found in Table S5.We used BWA 0.7.17, SAMtools1.9, and GATK 3.7 to generate genotype calls^80–83^. The pipelines are described in more detail in the next *Drosophila* section.

### Polymorphism Data

#### 1) Genus *Drosophila*

We downloaded Drosophila Population Genomics Project 3 consensus sequences from the Pool lab website (DPGP3)^84^. We obtained all 197 haploid embryo genomes collected from Zambia. *Drosophila simulan*s polymorphism data from 170 inbred individuals from an American population were obtained from Signor *et al*. (2018)^85^. Sequence data for 20 *D. yakuba* inbred lines are from Rogers *et al*. (2014)^86^. These lines are from Cameroon (10 lines) and Kenya (10 lines). *D. sechellia* data are from 41 outbred lines described in Schrider *et al*. (2018)^87^. These lines were collected from Praslin (7 lines), La Digue (7 lines), Marianne (2 lines), Mahé(7 lines), and Denis (18 lines) islands.

We aligned reads from each line using the BWA_MEM algorithm from BWA 0.7.17 to the appropriate reference sequences^80^. We then used SAMtools 1.9 to sort, index alignments, and pick properly paired alignments^81^. We used Picard (https://github.com/broadinstitute/picard) to mark and remove duplicates and performed local realignment with GATK’s RealignerTargetCreator and IndelRealigner tools (version 3.7)^83^. We used BCFtools mpileup with option “-Q 20 -q 20 -a DP, AD” and bcftools call with option “-cvO” for variant calling^82^. Multi-allelic sites and indels were excluded from analyses using VCFtools v0.1.17^88^option “--remove-indels --min-alleles 2 --max-alleles 2”.

#### 2) Genus *Arabidopsis* and *Capsella*

We used *A. thaliana* genome-wide polymorphism data sets from the 1001 Genomes Project^45,46^. Briefly, 1135 wild inbred lines, which represent the native Eurasian and North African range and recently colonized North America, were used for this study. Raw *Arabidopsis lyrata* sequence data were downloaded from NCBI and genotypes were called followed the protocol of Mattila *et al*. (2017)^89^. We downloaded raw *Arabidopsis halleri* reads from 54 individuals^90^. We obtained sequence data from 8 *Capsella grandiflora* samples from Josephs *et al*. (2015)^91^ and 12 *C. rubella* samples from Ågren *et al*. (2014)^92^. We used BWA 0.7.17^80^, SAMtools 1.9^81^, and GATK 3.7^83^ to generate genotype calls as described above for *Drosophila*.

#### 3) Order Primates

We downloaded genotypes of Han Chinese and Yoruba individuals as VCF files from the 1000 Genomes Project website^29,93^. We obtained genotypes of bonobos, chimpanzees, gorillas, and orangutans from Prado-Martinez *et al*. (2013)^32^ and the genome-wide polymorphism data set of four subspecies of Chinese rhesus macaques *Macaca mnulatta* from Liu *et al*. (2018)^94^. We obtained sequence data from 18 nub-nosed monkey *Rhinopithecus. roxellana* samples and 20 *R. bieti* samples from Yu *et al*. (2016)^74^. Finally, we downloaded eight old world monkey and one marmoset genomes (*Callithrix jacchus*) raw reads from NCBI (listed in Table S5). We again used BWA0.7.17^80^, SAMtools 1.9^81^, and GATK 3.7^83^ to generate genotype calls from the raw read data sets.

### Demographic history and Ka/Ks ratio dynamics

Based on the definition of *θ* (*θ* = 4Neμ), we estimate the long-term effective population size from *θ_W_* or *θ*π. Let *ξ_i_* be the number of synonymous segregating sites where the mutation occurs *i* times in the sample. We estimated Watterson’s *θ_W_* (1975)^95^, Tajima’s *θ_π_* (1983)^96^:

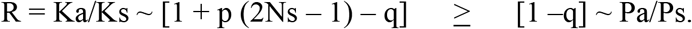

Where 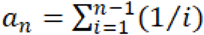.

We also inferred demographic history for humans, chimpanzees, and macaques using the pairwise sequentially Markovian coalescence (PSMC) model^28^. The PSMC human results were obtained directly from supporting data of the 1000 genomes projects^29^. We used the high coverage PCR-free sample (NA18525) from the Han Chinese population to represent *Homo sapiens*. The PSMC estimates from a Chinese rhesus macaques (CR1) were download from the supporting data in Liu *et al*. (2018)^94^. Chimpanzee genotypes from VCF files and callable regions BED files, download from the Great Ape Genome Project^32^, were transformed to the whole-genome diploid consensus sequence using VCFtools^88^. We then used “fq2psmcfa” to transform the consensus sequence into a fasta-like format where the i-th character in the output sequence indicates whether there is at least one heterozygote in a bin of 100 bp. Parameters were set as follows: ‘‘-N25 -t5 -r5 -p 4+25*2+4+6”. Human and chimpanzee generation time (g) was set at 20 years and the mutation rate (μ) at 1.25e-8 per site per generation^29,32^. Macaque generation time was set to 11 years and the mutation rate to 1e-8 per generation^94^.

In the last 5 million years, the lowest macaque effective population size was 22869 between 2.00 and 2.33 million years ago (see Fig. 1 and Table S6). When we assumed all non-synonymous mutations to be deleterious (q=1) with constant fitness effect of s = −2.186×10^−5^ (= −1/22869), the |Ns| of non-synonymous mutations is 0.04-0.62 in humans, 0.27-1.99 in chimpanzees, and 0.5-5.04 in macaques (see Table S6). The Eq (2’) was used to estimated R values in each time interval with known N and s. We estimate that R is 0.500-0.959 in humans, 0.076-0.757 in chimpanzee, and 4.2×10^−4^ – 0.582 in macaques.

### Theoretical Pa/Ps ratio spectrum

We assumed that all synonymous mutations are neutral and non-synonymous mutations either advantageous (p), deleterious (q), or neutral (1 – p - q). We also assumed a random mating population of constant size, infinite site model, free recombination, and no dominance (h = 0.5). s and s’ are the selection coefficients of advantageous and deleterious mutations, respectively. Let σ = Ns and σ’ = Ns’, the expected frequency distributions of advantageous or deleterious non-synonymous mutations^3,7,8,21^, be

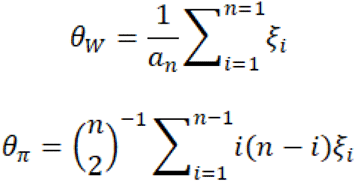

where x is the frequency of an advantageous polymorphism and θ_a_ = 4Nμ_a_ the number of new non-synonymous mutations in the population per generation. The expected frequency spectrum of the neutral non-synonymous polymorphism is (1 – p - q) θ_a_ / x and of the synonymous polymorphism is θ_s_ / x. Therefore, the Pa/Ps spectrum (θ_a_ = θ_s_) is

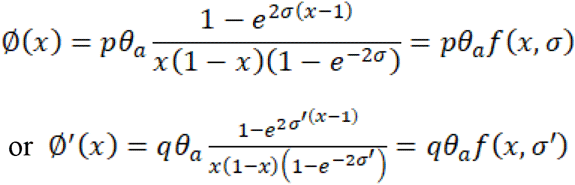

### Computation of Ka/Ks ratios and Pa/Ps ratios

We used an approximate method to estimate Ka/Ks ratio with an improved Nei-Gojobori model^97–99^. We estimated transition to transversion ratio (α/2β in Ina.1995^98^, or κ/2 in Yang el. 1998^100^) from the fourfold degenerate sites. Therefore, the proportion of transition mutations is κ / (2 + κ). We then used this ratio to count the expected number of synonymous and non-synonymous mutations in each codon. For example, the number of synonymous sites in TTT codon is S_TTT_ = 0 + 0 + κ / (2 + κ) because only the transition mutation at the third position of TTT is synonymous. Next, we counted the number of synonymous and nonsynonymous substitutions. When more than two differences exist between any two codons compared, we used the method described in Nei and Gojobori (1986)^97^. Finally, we used the Jukes-Cantor (1969)^101^ method to correct for multiple substitutions and obtain substitution rate estimates (Ka and Ks).

Because some genome assemblies used for polymorphisms calls have been updated, we changed the genome versions of some VCF files. For example, we downloaded the LiftOver file from UCSC and converted the genome coordinates of DPGP3 variants from dmel5 to dmel6 using CrossMap^102^(see Table S5). We then used SnpEff^103^ to annotate synonymous and non-synonymous changes for polymorphism based on the reference assembly. For *Drosophila* and *Arabidopsis* data, we used the free ratios model (model = 1, NSsites = 0, fix_omega = 0) of CODEML PAML module to infer ancestral sequences at each node on the known phylogenetic trees (see Fig. S1 and Fig. S2)^22,23^. The Ka/Ks ratios of each branch were then estimated using the method mentioned above. The allele identical corresponding to the ancestral branch was set as the inherited allele, and the alternative as derived. Thus, we were able to estimate the derived allele frequency. The computation of Pa/Ps ratios is similar to the method for the Ka/Ks ratio which is described above. We obtained Pa and Ps directly from observed polymorphism data and expected mutation counts without multiple hits correction.

## Acknowledgements

We thank Daniel Hartl, Ziheng Yang, Adam Eyre-Walker, Dan Graur, Justin Fay, Bruce Rannala and Matthew Hahn for their helpful discussions. This study was supported by the National Natural Science Foundation of China (31830005 and 31971540); the National Key Research and Development Plan (2017FY100705); the 985 Project (33000-18831107); and Guangdong Basic and Applied Basic Research Foundation (2019A1515010752).

