## Supplementary Information for "Two decades of suspect evidence for adaptive DNA-sequence evolution – Less negative selection misconstrued as positive selection"

**This file includes:**

Figures S1-S2

Tables S1-S2

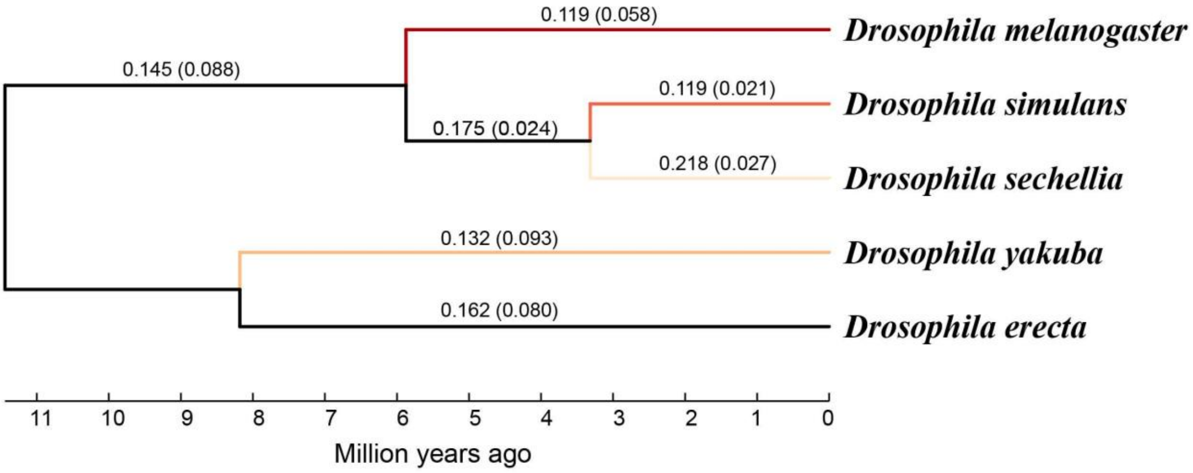


**Figure S1. Ka/Ks ratios (and Ks) along each branch of the *Drosophila melanogaster* subgroup phylogeny.** The Ka/Ks ratios of colored branches are the lineage-specific ratios shown in Fig. 3a.

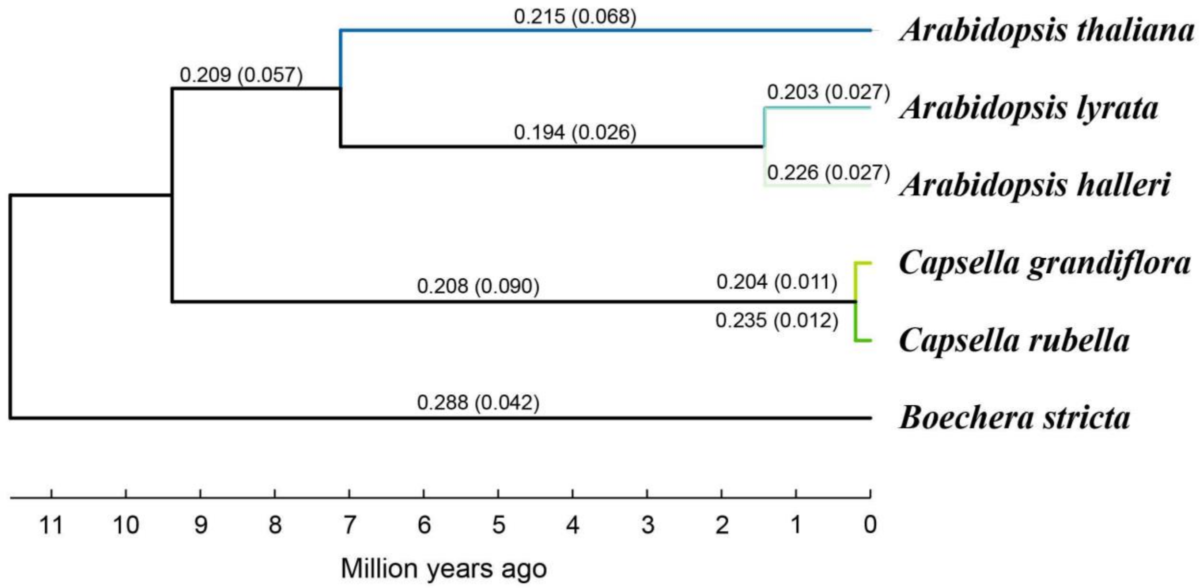


**Figure S2. Ka/Ks ratios (and Ks) along each branch of the *Arabidopsis thaliana* group phylogeny.** The Ka/Ks ratios of colored branches are the lineage-specific ratios shown in Fig. 3c.

**Table S1. Summary of non-synonymous and synonymous mutations in *Drosophila***

| | | ***D. melanogaster*** | ***D. simulans*** | ***D. sechellia*** | ***D. yakuba*** |
|---|---|---|---|---|---|
| **Sample Size** | | 197 | 170 | 41 | 20 |
| **$\theta\pi$ of synonymous polymorphisms ($\times 10^{-3}$)** | | 16.961 | 21.637 | 0.971 | 23.917 |
| **Expect non- synonymous sites** | | 7871815 | 7889518 | 7902719 | 7905260 |
| **Expect synonymous sites** | | 2884307 | 2866604 | 2853403 | 2850862 |
| **Polymorphism (all sites)** | Non-synonymous | 170387 | 53432 | 6673 | 79977 |
| | Synonymous | 379254 | 243919 | 10341 | 299429 |
| | Pa/Ps ratio | 0.165 | 0.080 | 0.233 | 0.096 |
| **Polymorphism (Freq. <= 0.2)** | Non-synonymous | 156274 | 34851 | 3022 | 65146 |
| | Synonymous | 297622 | 142543 | 4067 | 212080 |
| | Pa/Ps ratio | 0.192 | 0.089 | 0.268 | 0.111 |
| **Polymorphism (Freq. > 0.2)** | Non-synonymous | 14113 | 18581 | 3651 | 14831 |
| | Synonymous | 81632 | 101376 | 6274 | 87349 |
| | Pa/Ps ratio | 0.063 | 0.067 | 0.210 | 0.061 |
| **Lineage-specific Divergence** | Non-synonymous | 54366 | 19335 | 46372 | 96553 |
| | Synonymous | 161978 | 58365 | 75703 | 250424 |
| | Ka/Ks ratio | 0.119 | 0.119 | 0.218 | 0.132 |

**Table S2. Summary of non-synonymous and synonymous mutations in *Arabidopsis***

| | | *A. thaliana* | *A. lyrata petraea* | *A. lyrata lyrata* | *A. halleri* | *C. grandiflora* | *C. rubella* |
|---|---|---|---|---|---|---|---|
| **Sample Size** | | 1135 | 22 | 6 | 54 | 8 | 12 |
| **θπ of synonymous polymorphisms (×$10^{-3}$)** | | 5.645 | 15.885 | 2.986 | 7.082 | 18.451 | 3.179 |
| **Expect non- synonymous sites** | | 11661474 | 11607510 | 11604915 | 11619655 | 11638640 | 11618366 |
| **Expect synonymous sites** | | 4123581 | 4177545 | 4180140 | 4165400 | 4146415 | 4166689 |
| **Polymorphism (all sites)** | Non-synonymous | 356738 | 179687 | 26449 | 147021 | 137199 | 27699 |
| | Synonymous | 314737 | 250125 | 32016 | 143511 | 252799 | 44925 |
| | Pa/Ps ratio | 0.401 | 0.259 | 0.298 | 0.367 | 0.193 | 0.221 |
| **Polymorphism (Freq. <= 0.2)** | Non-synonymous | 327471 | 116453 | 8331 | 109754 | 92545 | 17132 |
| | Synonymous | 252773 | 128809 | 7895 | 85201 | 142929 | 25370 |
| | Pa/Ps ratio | 0.458 | 0.325 | 0.380 | 0.462 | 0.231 | 0.242 |
| **Polymorphism (Freq. > 0.2)** | Non-synonymous | 29267 | 63234 | 18118 | 37267 | 44654 | 10567 |
| | Synonymous | 61964 | 121316 | 24121 | 58310 | 109870 | 19555 |
| | Pa/Ps ratio | 0.167 | 0.188 | 0.271 | 0.229 | 0.145 | 0.194 |
| **Lineage-specific Divergence** | Non-synonymous | 168096 | 58212 | 64210 | 69658 | 27008 | 31663 |
| | Synonymous | 266642 | 105259 | 112341 | 109138 | 46955 | 48105 |
| | Ka/Ks ratio | 0.215 | 0.196 | 0.203 | 0.226 | 0.204 | 0.235 |
